# Eye movements modulate neural activity in the human anterior thalamus during visual active sensing

**DOI:** 10.1101/2020.03.30.015628

**Authors:** Marcin Leszczynski, Tobias Staudigl, Leila Chaieb, Simon Jonas Enkirch, Juergen Fell, Charles E. Schroeder

## Abstract

Humans and other primates explore visual scenes by *active sensing*, using saccadic eye movements to relocate the fovea and sample different bits of information multiple times per second. Saccades induce a phase reset of ongoing neuronal oscillations in primary and higher-order visual cortices and medial temporal lobe. As a result, neuron ensembles are shifted to a common state at the time visual input propagates through the system (i.e., just after fixation). The extent of the brain’s circuitry modulated by saccades is not yet known. Here, we evaluate the possibility that saccadic phase reset impacts the anterior nuclei of the thalamus (ANT). Using rare recordings in the human thalamus of three surgical patients, we found saccade-related phase concentration, peaking at 3-4 Hz, coincident with suppression of Broadband High-frequency Activity (BHA; 80-180 Hz). Our results provide evidence for saccade-related modulation of neuronal excitability dynamics in the ANT, consistent with the idea that these nuclei are engaged during visual active sensing. These findings show that during real-world active visual exploration neural dynamics in the human ANT, a part of extended hippocampal–diencephalic system for episodic memory, exhibit modulations that might be underestimated in typical passive viewing.

## Introduction

Both human and non-human primates sample visual scenes actively by systematically shifting eye gaze several times per second (Yarbus, 1967; Gilchrist et al., 1997). These eye movements modulate neural activity along the visual pathway from the lateral geniculate nucleus (e.g., Lee and Malpeli, 1998; Reppas et al; 2002; Sylvester et al., 2005; Sylvester & Rees, 2006; McFarland et al., 2015) through ascending stages of visual cortex (Purpura et al., 2003; Rajkai et al., 2008; Bartlett et al., 2011; Ito et al., 2011; Hamamé et al., 2014; Zanos et al., 2015; 2016; Barczak et al., 2019) up to the medial temporal lobe (MTL) including the hippocampus and entorhinal cortex (Killian et al., 2012; Hoffman et al., 2013; Jutras et al., 2013; Killian et al., 2015; Staudigl et al., 2017; Meister & Buffalo, 2018; Staudigl et al., 2018; Katz et al., 2018; Doucet et al., 2019; for a recent review see Leszczynski and Schroeder, 2019).

Earlier studies of the effects of saccades in the dark (Ringo et al., 1994; Sobotka & Ringo, 1997; Lee and Malpeli, 1998; Sylvester et al., 2005; Sylvester & Rees, 2006; Royal et al., 2006; Rajkai et al., 2008; Kagan et al., 2008; Nakamura & Colby, 2000), and more recent studies that minimize saccade-related visual input by various means (McFarland et al., 2015; Barczak et al., 2019) confirm the proposition that nonretinal “corollary discharge” (CD) signals generated in parallel to saccades, modulate the excitability of neurons in visual pathways structures (Sommer & Wurtz, 2008). Most recently, Barczak et al. (2019) showed that nonretinal saccadic signals reset ongoing excitability fluctuations (oscillations) in V1 neuron ensembles to a high excitability phase, and that this effect amplifies their response to visual inputs arriving immediately after the saccade (i.e., at fixation onset). This phase modulation is primarily observed in theta and alpha ranges - key physiological signatures of active sensing (Ahissar and Arieli, 2001; Szwed et al., 2003; Kleinfeld et al., 2006; Leszczynski and Schroeder, 2019).

Interestingly, many of the areas displaying saccadic phase modulation lie outside of the classic visual system - higher order regions with multiple sensory and cognitive functions like the MTL. However, aside from the tectal nuclei (e.g., Lee et al., 1988; Munoz & Wurtz, 1995; Werner-Reiss & Groh et al; Sommer & Wurtz, 2008) and the lateral geniculate nucleus of the thalamus (Lee and Malpeli, 1998; Reppas et al., 2002; Sylvester et al., 2005; Sylvester & Rees, 2006; Royal et al., 2006; McFarland et al., 2015), the degree to which saccade-related signals impact other subcortical structures has not been systematically investigated. As a step toward addressing this question, we leveraged a unique dataset: intracranial electroencephalography (iEEG) recordings from anterior nuclei of the thalamus (ANT) in patients implanted for deep brain stimulation treatment of epilepsy (see e.g. Lehtimäki et al. 2018; see Figure 1A).

**Figure 1.**
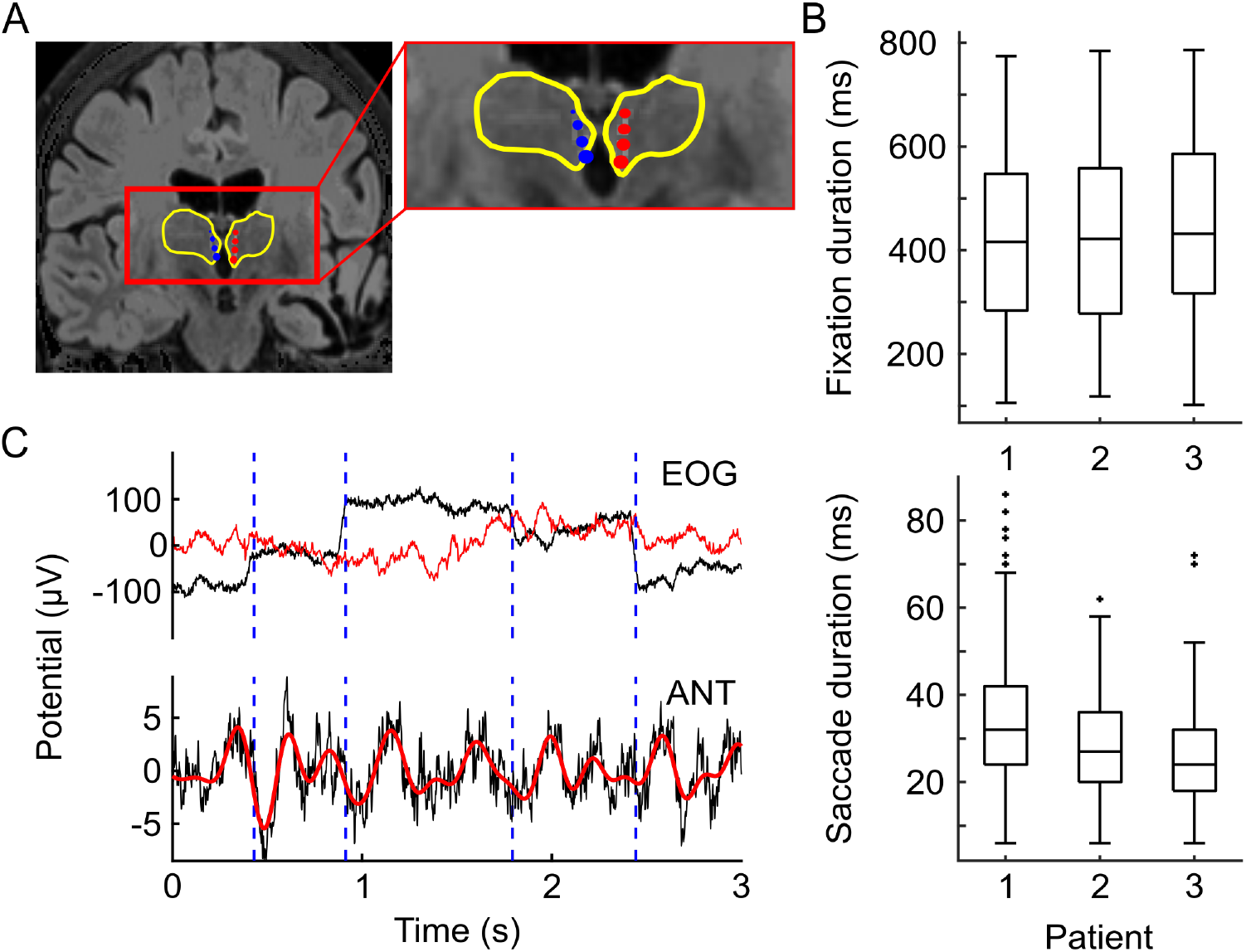
Implantation trajectory, example field potential trace and behavior. **(A)** Bilateral leads implanted into the anterior nuclei of the thalamus marked with blue (right ANT) and red (left ANT) in a single patient (4 contacts per side). Inter-contact space of 3 mm (contact height of 1.5 mm with 1.5 mm spacing). Yellow contours depict boundaries of the thalamus. **(B)** Distribution of fixation (upper panel) and saccade (lower pannel) durations for three patients. Box plots indicate 25^th^, median and 75^th^ percentile, whiskers extend to extreme values not considered outliers while outliers are marked with crosses. Because we used the EOG signal to define time points of fixation and saccade onsets, it is possible that smaller eye movements were not detected (see Limitations). **(C)** An example of 3 seconds long recording trace with signals from EOG (black trace) and Fp1 (red trace). Horizontal dashed lines indicate fixation onset time points (upper panel). The same interval of 3 seconds field potentials from an example site in the ANT. Black trace presents raw signal; red trace shows signal filtered in 2-5 Hz (lower panel).

Neural activity can be enhanced or suppressed by the specific phases of ongoing oscillations (Lakatos et al., 2005; Mormann et al., 2005; Canolty et al., 2006; see also Leszczynski et al., 2017). This suggests that neural response should be increased or decreased by resetting local neural oscillations depending on the exact phase to which the oscillation is reset to. Indeed, previous studies showed that neural oscillations might be reset to a high-excitability phase (i.e., the one that during which neural activity is enhanced) but also to a low-excitability phase (i.e., during which neural activity is attenuated). Recordings from primary visual cortex (V1) showed that attention to one out of two interleaved stimulus streams (visual or auditory) entrains supragranular layers delta oscillation with high-excitability phases aligned to the attended stimulus stream and low-excitability phases aligned to the unattended stream (Lakatos et al., 2008). These two attention states were associated with counter-phase modulations of neural activity. Increased current source density (CSD), multi-unit activity (MUA) and BHA magnitudes occurring around the negative peak (high-excitability phase), while the smallest response amplitudes occur around the positive peak (low-excitability phase) of delta oscillations right before the stimulus presentation (Lakatos et al., 2008). Similar effects were observed in the primary auditory cortex (A1) - somatosensory stimuli reset neural oscillations to different phase angles with distinct effects of ipsi- and contra-lateral somatosensory stimulation on the supra- and infragranular layers MUA in A1 (Lakatos et al., 2007; see also Lakatos et al., 2009 for similar effects between A1 and V1). Altogether, these findings show a supramodal mechanism by which attention can control neural oscillations by resetting them to either high- or low- excitability phase. While previous studies on visual active sensing showed that phases across the visual system are reset to a high-excitability state it is also possible that neural activity outside of the visual system is reset to a low-excitability state. This would be evident as an increased phase coherence coupled with a decrease in BHA at fixation onset.

Based on the interconnectivity of the ANT with other brain regions like the hippocampus and anterior cingulate cortex (Bubb et al., 2017; Zikopoulos and Barbas, 2006) as well as their apparent receipt of direct retinal afferents (Itaya et al., 1986) we hypothesized that neuronal excitability in the ANT should be modulated by saccade-related signals. We tested three specific predictions: **1)** Saccades should perturb ongoing field potential activity in the human ANT; **2)** The phase of low frequency oscillations should cluster at or just after fixation onset; **3)** Broadband High-frequency Activity (BHA; 80-180 Hz), a measure of neuronal processes correlated with neuronal spiking (Mukamel et al., 2005; Niessing et al., 2005; Nir et al., 2007; Ray et al., 2008; Leszczynski et al., 2020), should be systematically modulated by the saccade-fixation cycle - an increase would suggest reset to a high- while a decrease would suggest reset to a low-excitability phase. Our findings provide support for all three hypotheses and show that the ANT is reset to a low-excitability phase. Thus, in addition to a range of other functions, the ANT appears to be engaged during visual active sensing.

## Results

### Fixation-locked field potentials in human ANT

Using frontal scalp EEG and EOG electrodes we detected time points of saccade and fixation onset (Figure 1B–C; see Methods). This is possible because eye movements generate large magnitude electric field fluctuations that are measurable at scalp EEG across multiple frontal channels (Lins et al., 1993). Combining simultaneous EEG and direct ANT recordings we could thus study the impact of eye movements (as detected from EEG) on neural activity in human anterior nuclei of the thalamus.

As in our earlier studies (Rajkai et al., 2008; Barczak et al., 2019), we focus on neural events related to the end of the saccade (fixation onset) as this is an event with clear perceptual relevance; i.e., the point at which retinal inflow begins in the ascending visual pathways. First, we tested whether fixation onset-related potentials (FRPs) are modulated relative to a surrogate distribution. The surrogates (N=1000) were created by locking intact segments of ANT field potentials to pseudo-fixations (i.e., random time points uniformly distributed across the entire recording session; see Methods). We observed that the magnitude of fixation onset-related potentials departs from the surrogate distribution at multiple time points (permutation test; p < 0.05; FDR controlled for multiple comparisons; see Methods). An initial positive deflection around 100 ms before fixation onset was followed by a negative deflection peaking at 40-50 ms after fixation onset and another positive deflection peaking around 160 ms after fixation onset (Figure 2A). We also observed that FRPs exceeded the surrogate distribution in each of the three participants (see Figure 2E,I,M).

**Figure 2.**
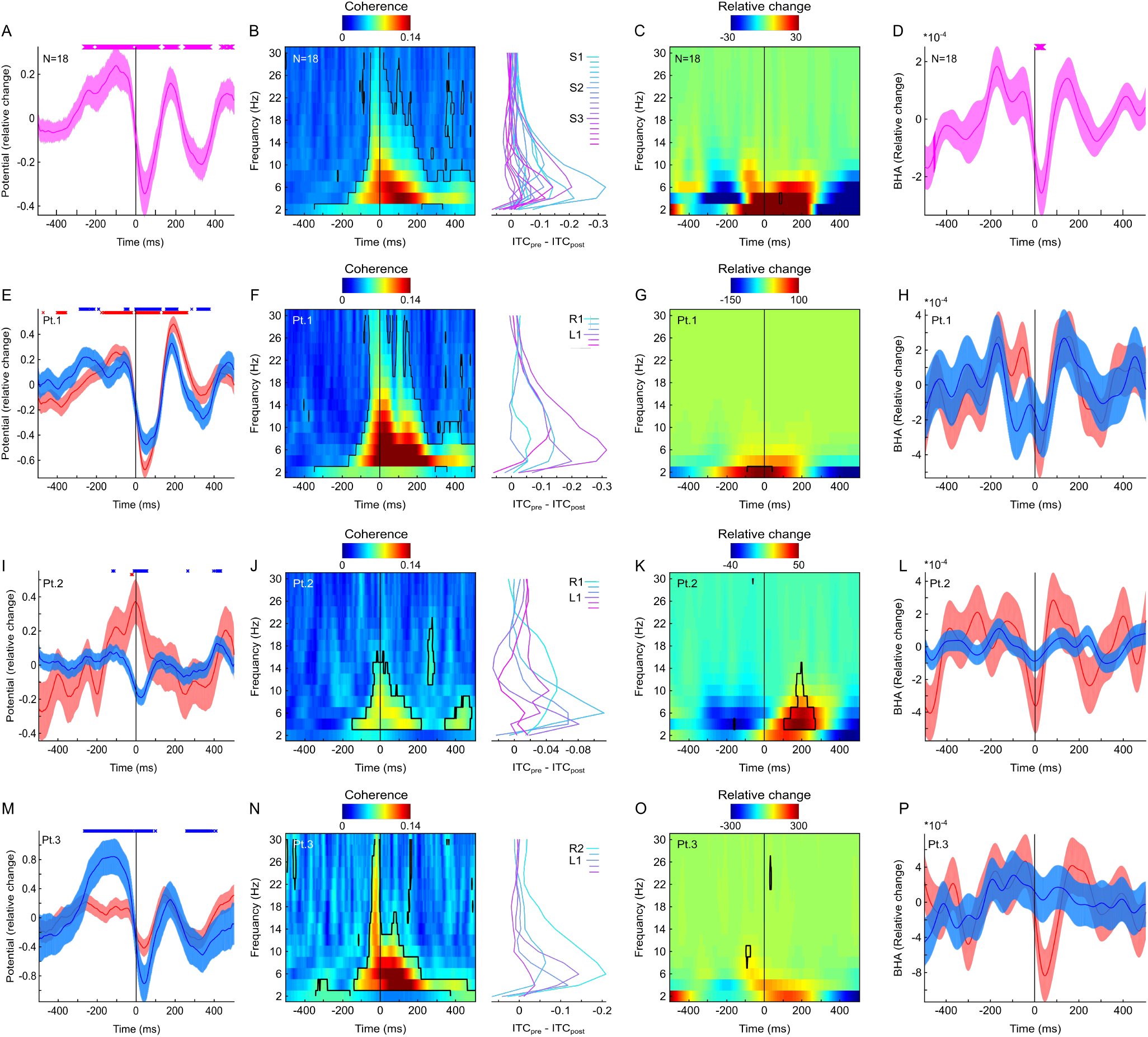
Fixation-locked neural activity in human ANT. **(A)** Grand average fixation-locked field potentials (N = 18 ANT channels, 3 patients). Shading reflects standard error of the mean (SEM). Vertical lines show the point of fixation onset. Markers above indicate significant time points (p < 0.05; permutation test). **(B)** Color map shows inter-fixation phase coherence (time on x-axis, frequency on y-axis). Contours depict significant time-frequency points (permutation test p < 0.05). Vertical black line indicates fixation onset. Line plots on the right side indicate ITC change from pre- (−300:−100 ms) to post-fixation (0:100 ms) time window separately for each channel. **(C)** Color map shows fixation locked power relative to trial average (time on x-axis, frequency on y-axis). Contours present change from surrogate distribution (permutation test p < 0.05). **(D)** Fixation-locked BHA (80-180Hz). Shading reflects SEM. Vertical black line indicates fixation onset. Markers above indicate significant time points (permutation test p < 0.05). **(E)** Fixation-locked field potentials (patient 1) from channels on the right ANT (blue) and left ANT (red). Shading reflects standard error of the mean (SEM). Vertical black lines show the point of fixation onset. Markers above indicate significant time points (permutation test p < 0.05). **(F)** Same as B but for data from patient 1. **(G)** Same as C but for data from patient 1. **(H)** Fixation-locked BHA (patient 1) from channels on the right ANT (blue) and left ANT (red). Shading reflects standard error of the mean (SEM). Vertical black lines show the point of fixation onset. Data from patient 2 **(I-L)** and patient 3 **(M-P)** plotted with the same convention as (E-H). Note that the reversed polarity in patient 2 left channels might be explained by a more posterior and dorsal trajectory of the left electrode shaft. Furthermore, patient 3 had the right side electrode placed deep with ventral channels showing no signal (note only R2 and R3 are presented; see Methods for more details). All results are controlled for multiple comparisons with Benjamini & Hochberg/Yekutieli false discovery rate procedure.

### Fixation-locked low frequency phase, power and BHA

Next, we investigated how the saccade-fixation cycle influences spectral phase and power in the ANT. To this end, we studied inter-fixation phase coherence, perisaccadic power spectra across a range of low frequencies (1-30 Hz) and BHA power (80-180 Hz; see Methods). Overall, we found that fixation-locked phase coherence increased above the 95^th^ surrogate percentile (corresponding to p < 0.05; FDR controlled for multiple comparisons) across multiple time and frequency points with strongest effects of phase clustering at 3-4 Hz around 120 ms after fixation onset (see Figure 2B). This effect was evident in each of the three patients (see Figure 2F,J,N). In contrast we detected a weak modulation of fixation-locked spectral power relative to their surrogate distributions (Figure 2C) which was also weak and unreliable at each of the patients when quantified separately (Figure 2G,K,O). We found a small fixation-locked BHA decrease below the 5^th^ percentile of the surrogate distribution around 40-50 ms after fixation onset (Figure 2D). This effect was small but reliable at the aggregated level (Figure 2D). While it did not exceed single subject surrogate distribution it showed consistent direction with the dip in the BHA observed right after fixation onset for each of the patients (Figure 2H,L,P).

### Distinct modulation of the ANT field potentials by eye blinks

Next, we tested the influence of eye blinks on the ANT field potentials. We observed a negative ERP deflection during the eye blink which peaked about 50-60 ms before the blink offset. It was followed by a positive deflection peaking about 100-140 ms after blink offset. We also found ITC exceeding the surrogate level across multiple time and frequency points with the strongest effects visible in frequencies below 10 Hz. We observed weak to undetected power modulations. Finally, the BHA showed a weak decrease during the blink which did not survive correction for multiple comparisons and a rebound that peaked about 200 ms after the blink offset which is similar to previous findings on the BHA modulations by the eye blinks (Golan et al., 2016). Fixation onset and eye blink offset both mark the time points of changes to the retinal input. Despite this, there are few important differences between ANT field potentials locked to these two eye events suggesting that the neural dynamics in the ANT reflects something else than a change to retinal input. First, ERPs locked to fixation onset and eye blink offset show different morphology. While fixation-locked ERPs appear to oscillate for at least 2 cycles, eye blink offset locked ERPs are transient and return to baseline right after the blink offset. The ITC shows similar spectral but distinct temporal profiles. Fixation-locked ITC is strongest after the event. The blink offset-locked ITC is more uniformly distributed before and after the eye event. Finally, the BHA shows a transient decrease right after fixation onset. In turn, the blink offset locked BHA shows an increase peaking about 200 ms after the event which is similar to previous observations (Golan et al., 2016). Altogether, these results show that the ANT is modulated by both saccade-fixation cycle as well as the eye blinks but the morphology of these effects is distinct.

### Potential limitations of this study

We analyzed data from only three patients. However, this in itself is not a critical limitation, as we have a large number of repeated measures within the individual patients (fixations: N = 1019, 820, 637) and our results are visible at multiple ANT channels and reproduced in each participant. A more obvious limitation is that while EOG-based eye movement detection provides great temporal precision and high accuracy in detecting large saccades comparable with modern eye tracking (Lins et al., 1993; Joyce et al., 2002; Toivanen et al., 2015; Mueller et al., 2016; Jia & Tyler, 2019), it is less sensitive to small eye movements. It is therefore possible that the current study underestimates the contribution of shorter saccades (and microsaccades) and over-estimates fixation durations. To ensure that our results are robust across fixation durations, we performed control analyses in which we limited maximum duration of fixation to 600 and 800 ms resulting in distributions with median durations centered at 300 ms or 500 ms (comparable to those known in real life visual exploration; e.g., Hayhoe et al., 2003). Both of these control analyses showed similar results – significant ERP deflection, increased phase coherence, decreased BHA and no weak to undetected effects on power. The EOG-based eye tracking provides little information about the direction of gaze. Thus, while reporting saccadic modulation of ANT activity, our study leaves open the question of whether gaze direction is systematically reflected in the field potentials of ANT activity. We used Fp1 referenced to linked mastoids for detecting vertical part of EOG in patients 1-3 and bipolar EOG 2 - EOG1 in patients 1 and 3 and F7 – F8 in patient 2 for detecting horizontal EOG. EOG signals have been shown to spread across all these channels with varying magnitude (Lins et al., 1993). Despite these slight differences in electrode placement, we find consistent effects and similar eye movement distributions across all three patients, confirming that our results are robust across these variations in electrode position. Overall, there is no reason to think these limitations have a major influence on our main conclusions. Finally, previous studies recorded eye movements in dark to separate stimulus elicited and saccade elicited signals. However, while recording in the dark is critical to isolate influence of eye movements on the neural dynamics from the visual evoked potential in the visual system (where eye-locked perturbations and visual evoked potential temporally overlap), it is less critical outside of the visual system, where these two signal might be temporarily separated. There are at least two reasons to think that the current results reflect eye movement related signals rather than visual evoked potential. First, previous study on human ANT showed that the visual evoked potential in the ANT peaks around 300 ms after stimulus onset with earliest measured latency of about 240 ms post stimulus followed by a slow negative potential peaking around 700-1000 ms after stimulus onset (Štillová et al., 2015). While we also see this P300-like component in the current data, the main ERP deflection is much faster (<100 ms) suggesting that eye movement and visual evoked potential might have distinct temporal profiles in the human ANT. This is further supported by comparison of fixation onset and eye blink offset locked analyses. Both fixation onset and eye blink offset mark the time point of change to the retinal stimulation. Thus, if the current results reflect changes to the visual input, we should observe little to no difference between field potentials following saccade and eye blink offset. Because the morphology of the ERP, ITC and BHA are different for fixation-onset and blink-offset (compare Figure 2 and Figure 3), we reasoned that these effects reflect something else than visual evoked potential.

**Figure 3.**
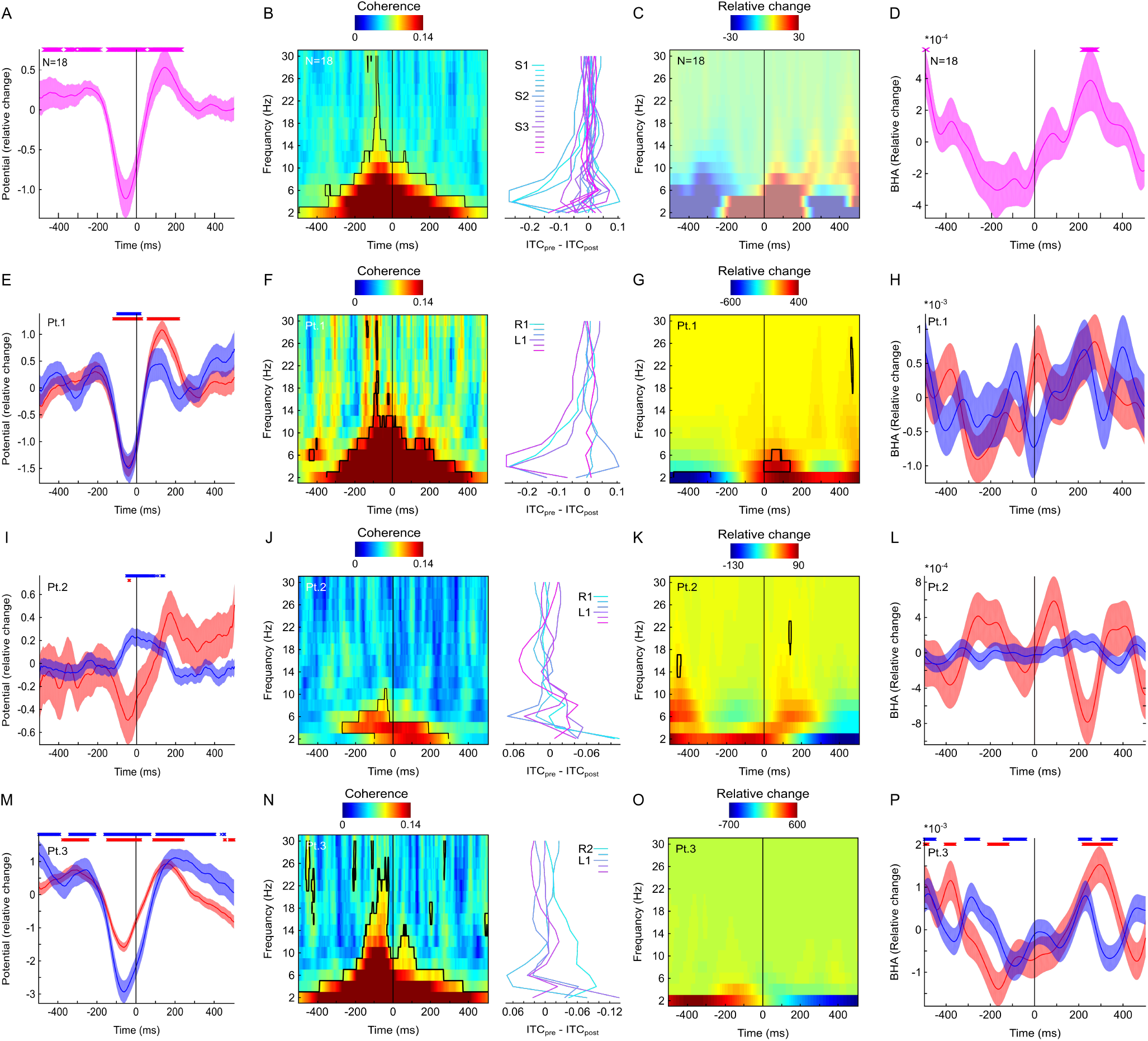
Blink offset-locked neural activity in human ANT. **(A)** Grand average for blink offset-locked field potentials (N = 18 ANT channels, 3 patients). Shading reflects standard error of the mean (SEM). Vertical lines show the point of blink offset (i.e., onset of visual input). Markers above indicate significant time points (permutation test p < 0.05). **(B)** Color map shows inter-fixation phase coherence (time on x-axis, frequency on y-axis). Contours depict significant time-frequency points (permutation test p < 0.05). Vertical black line indicates blink offset (i.e., the beginning of visual input). Line plots on the right side indicate ITC change from pre- (−300:−100 ms) to post- blink offset (0:100 ms) time window separately for each channel. **(C)** Color map shows blink offset locked power relative to trial average (time on x-axis, frequency on y-axis). No change from surrogate distribution (permutation test p < 0.05). **(D)** Blink offset locked BHA (80-180Hz). Shading reflects SEM. Vertical black line indicates blink offset (i.e., beginning of visual input). Markers above indicate significant time points (permutation test p < 0.05). **(E)** Blink offset-locked field potentials (patient 1) from channels on the right ANT (blue) and left ANT (red). Shading reflects standard error of the mean (SEM). Vertical black lines show the point of blink offset (i.e., beginning of visual input). Markers above indicate significant time points (permutation test p < 0.05). **(F)** Same as B but data from patient 1. **(G)** Same as C but data from patient 1. Contours present change from surrogate distribution (permutation test p < 0.05) **(H)** Blink offset-locked BHA (patient 1) on the right ANT (blue) and left ANT (red). Shading reflects standard error of the mean (SEM). Vertical black lines show the point of blink offset. Data from patient 2 **(I-L)** and patient 3 **(M-P)** plotted with the same convention as (E-H). All results are controlled for multiple comparisons with Benjamini & Hochberg/Yekutieli false discovery rate procedure.

## Discussion

To evaluate the possible role of the anterior nuclei of the thalamus in visual active sensing, we studied neural dynamics surrounding fixations and eye blinks during unconstrained free viewing, in three epilepsy patients implanted with ANT electrodes for clinical purposes. Comparing pre- and post-fixation measures to surrogate distribution, we found significant fixation-related increases in field potential magnitude and phase coherence, albeit with weak to undetectable concomitant changes in low frequency power. Fixation-related BHA, a measure of neuronal processes correlated with neuronal population spiking (Mukamel et al., 2005; Niessing et al., 2005; Nir et al., 2007; Ray et al., 2008; Leszczynski et al., 2020), showed decrease from the surrogate distribution within the first 50 ms after fixation onset. This suggests that neural activity is suppressed in the ANT right after fixation onset. We also observed an ERP locked to fixation-onset with negative deflection peaking about 50 ms post-fixation, followed by a positive deflection about 200 ms after fixation and another negativity at about 300 ms after fixation onset. The biggest ERP component observed in the current study (i.e., N50) appears much faster than stimulus-locked evoked potentials in the ANT observed previously in response to passive viewing of visual stimuli (Štillová et al., 2015) in which the authors observed P300 potential with the shortest reported latency of about 240 ms. In their report the visual evoked P300 component was followed by a slow negative potential peaking around 700-1000 ms after stimulus onset. The distinct temporal profile of fixation-locked ERPs from that observed in passive viewing suggests that the ANT is differently involved during active and passive vision. This is further supported by our blink offset-locked analyses which show distinct ERP, ITC and BHA morphologies.

Fixation-related phase concentration was strongest around the rate of saccades (3-4 Hz), peaking about 120 ms after fixation onset. The impact of saccades on oscillatory phase coherence in the ANT is consistent with findings from the MTL (Killian et al., 2012; Hoffman et al., 2013; Jutras et al., 2013; Killian et al., 2015; Staudigl et al., 2017; Meister & Buffalo, 2018; Staudigl et al., 2018; Katz et al., 2018; Doucet et al., 2019) and the visual system (Purpura et al., 2003; Rajkai et al., 2008; Bartlett et al., 2011; Ito et al., 2011; Hamamé et al., 2014; Zanos et al., 2015; 2016; Barczak et al., 2019). However, the dynamics of fixation-related excitability modulation in the ANT, with a decrease in BHA around fixation onset, appears distinct from those reported for V1 (see e.g., Rajkai et al., 2008; Ito et al., 2011; Barczak et al., 2019). For example, Rajkai et al., (2008) observed the strongest suppression of neuronal firing during saccade and before the onset of fixation, and essentially the same thing was observed by Barczak et al., (2019). Interestingly, dynamics similar to those in the ANT have been noted in higher order visual areas. For example, Zanos et al., (2016) observed bimodal temporal distribution of suppression in V4 neurons, with late suppression being strongest after fixation onset. The decrease in BHA right after fixation onset coupled with strong increase in phase coherence observed in the current study, suggests that the ANT field potentials might be reorganized to a low-excitability state.

What function might the ANT play in visual active sensing? The anatomical connections of the ANT provide some clues as to its position in the visual active sensing network. The ANT receive descending inputs from the subiculum and ascending inputs from the mammillary bodies forming a central component of the extended hippocampal–diencephalic system for episodic memory (Vann and Aggleton, 2004). The ANT also has dense reciprocal connections to cingulate areas including the anterior cingulate cortex, which are implicated in top-down cognitive control (for review see Bubb et al., 2017). Studies in non-human primates (Itaya et al., 1986) as well as tree shrews and rats (Conrad and Stumpf, 1975; Itaya et al. 1981) have also identified direct input pathways from the retina and pretectum (Sikes and Vogt, 1987) to the anterodorsal thalamus. Studies in nonhuman primates (Zikopoulos & Barbas, 2006) demonstrate that anterior thalamic nuclei have reciprocal connections to frontal and prefrontal areas, and through their intrathalamic connections, indirect access to posterior visual areas, placing the ANT within the circuits that could mediate top-down modulation due to both attention and eye movements.

Functional properties of ANT neurons are consistent with a role in the organization of sensing activities. In rodents, neurons in the ANT fire as a function of the animal’s head direction (Taube, 1995; Tsanov et al., 2011; for review see Jankowski et al., 2013), providing crucial input during spatial navigation (Winter et al., 2015) by encoding animal’s perceived directional heading relative to its environment (Taube, 2007). This is important to the present discussion, as rodents tend to favor head movements over saccades in active sensing. Although vestibular inputs are critical for generating head direction firing, the directional signal appears to also involve the motor system (Taube, 2007). Taube & Burton (1995) showed that preferred direction firing is stable as the animal “actively” moves from one environment to another. However, “passive” transportation of an animal between two environments distorts directional firing and this distortion seems to be independent of the visual input (Stackman et al. 2003; for review see Taube, 2007). Overall then, the interconnectivity patterns of the ANT together with the functional properties of ANT neurons suggest that the ANT contributes more to top-down control, than to bottom-up input processing in active sensing.

How does mediation of control functions in active sensing fit with other functions that have been attributed to the ANT? The ANT have been proposed to contribute to a range of cognitive functions including learning and memory (Aggleton and Brown, 1999; Warburton et al., 2001; Vann and Aggleton, 2004) as well as attention (Bourbon-Teles et al., 2014; Wright et al., 2015; see also Leszczynski and Staudigl, 2016). Damage to the ANT or its inputs from the mammillary bodies leads to episodic memory deficit observed in Wernicke-Korsakoff and thalamic stroke patients (Kerrigan et al., 2004). Interestingly, along with other cognitive deficits, both Wernicke-Korsakoff and thalamic stroke patients have difficulty in motor sequence programming (Ghika-Schmid and Bogousslavsky, 2000; see also Carrera and Bogousslavsky, 2006). This last may be a key observation, because overt active sensing depends on motor sampling routines - sequences of movements by the eyes, whiskers, head, hands and breathing musculature. In an active sensing framework, these motor sequences imbue individual’s sampling strategy with momentary predictions and allow to transcribe the sensory world into a set of neural representations that the brain can use to generate perception and to guide further adaptive behavioral routines (Ahissar and Arieli, 2001; Szwed et al., 2003; Kleinfeld et al., 2006; Schroeder et al., 2010; Leszczynski and Schroeder, 2019).

## Methods

### Participants

Depth field potentials along with the surface EEG were recorded from 3 pharmacoresistant epilepsy patients (age range from 22 to 52, 2 male and 1 female) implanted with electrodes targeted to the anterior nucleus of the thalamus for treatment of epilepsy. Recordings were performed at the Department of Epileptology, University of Bonn, Germany. The study was approved by the Ethics Committee of the University of Bonn Medical Center. All the methods were performed in accordance with the relevant guidelines and regulations and all patients gave written informed consent.

### Electrophysiological recordings

The data were recorded using bilaterally implanted multi-contact (4 channels per shaft) depth electrodes and simultaneous scalp surface EEG electrodes (Fp1, Fp2, F3, F4, C3, C4, ′P3, P4, O1, O2, F7, F8, T3, T4, T5, T6, Cz, Fz, Pz, T1, T2, Cb1, Cb2) in patients 2 and additionally EOG1, EOG2 in patients 1 and 3 placed according to the 10-20 system. The depth electrodes’ contacts (Medtronic 3387) were of 1.5 mm height with an inter-contact center-to-center spacing of 3 mm. All data were sampled at 1kHz, on-line band pass filtered between 0.01Hz (6dB/octave) and 300Hz (6dB/octave), off-line down-sampled to 500Hz. The ANT depth channels were re-referenced to its neighbor using bipolar montage to increase local specificity of the signal. This results in 3 bipolar pairs per subject per side. In total we analyzed 18 bipolar pairs (3 pairs on shaft × 2 sides × 3 subjects). The data consist of continuous 1 hour long unconstrained recordings during which participants performed various cognitive tasks involving presentation of visual stimuli on a laptop computer, interacted with experimentators and members of the hospital staff.

### Electrode localization

Surgical procedures followed these described in the paper by Lehtimäki et al., 2018. Briefly, the locations of contacts were estimated relative to midcommissural point (MCP) using the AC-PC coordinate system and relative to visible landmarks such as mamillothalamic tract (MTT) ANT junction (MTT/ANT junction). The procedure utilizes both visible anatomical landmarks in individual patient MRIs and stereotactic atlas information. The most relevant anatomical landmark is the MTT that joins ANT nucleus at its inferior border slightly anterior to the midpoint of ANT in the anterior–posterior axis.

Preoperative navigation sequences were acquired with a 3.0-T MR imaging unit (Achieva, Philips Healthcare, Best, the Netherlands) and included high-spatial-resolution three-dimensional T1-weighted sequences pre- and postcontrast (isotropic three-dimensional gradient echo; voxel size, 1.0 × 1.0 × 1.0 mm; echo time, 3.48 msec; repetition time, 7.53 msec). Correct electrode placement was verified by postoperative high-spatial-resolution CT (Brilliance 16, Philips Healthcare, Best, the Netherlands; voxel size, 0.8 × 0.8 × 0.8 mm).

The electrode were first mapped onto brain using co-registration by iELVis (Groppe et al., 2017) followed by electrode identification on post-implantation CT co-registered to pre-implantation T1 image. To obtain the anatomical location labels presented in Figure 1, we used Freesurfer’s automated segmentation (Fischl et al., 2002).

In the current study patient 3 had the right side electrode placed deep with ventral channels showing no signal (note only R2 and R3 are presented in Figure 2 and 3). Furthermore, careful inspection of ERPs on the left and right ANT shaft suggests that right side ERPs in patient 2 are consistent with other patients, while the ERPs recorded from the left shaft show reversed ERP polarity (see Figure 2). This difference might be explained by a more posterior trajectory of the left ANT electrode with the leads locations being both posterior more dorsal compared to that on the right side. This suggests that the channels on the left side might be picking up signals from another nuclei which would explain ERP polarity reversal.

### Data analyses

All data were processed offline using Fieldtrip toolbox (Oostenveld et al., 2011) and MATLAB (MathWorks). Pre-processing was performed at single electrode level. Continuous data were down-sampled to 500Hz. To remove line noise we used a butterworth filter (4th order) at 50 Hz, 100 Hz, 150 Hz. Next, we created bipolar montage to maximize spatial specificity of the signal and also to decrease common noise and contributions from distant sources through volume conduction. To this end, we subtracted signals from neighboring contacts on the electrode (i.e., ANTR0 – ANTR1, ANTR1 ANTR2, ANTR2 – ANTR3 for right side contacts and ANTL0 – ANTL1, ANTL1 – ANTL2, ANTL2 – ANTL3 for left side contacts, respectively). ANTR0 is the deepest contact on the right shaft. Next, to calculate Broadband High-frequency Activity (BHA), we band-pass filtered (70 - 150 Hz) and hilbert-transformed continuous data. The complex-valued signal was then rectified and squared to extract power of the BHA. Low frequency activity (1-30Hz) was extracted from continuous data with 3 cycles Wavelets.

Next, we used scalp channels to create continuous signal maximizing vertical and horizontal components of the Electrooculogram (EOG). The EOG is a technique for measuring corneo-retinal standing potential that exists between the front and the back of the human eye. Several previous reports established the feasibility of measuring eye movements with this method. We acknowledge the limited spatial specificity of the EOG-based eye tacking in the limitations section and withhold from any statements regarding gaze location or direction of movement. At the same time EOG has excellent (i.e., comparable to modern eye tracking temporal resolution which makes it suitable for the purpose of detecting time points of fixations, saccades and eye blinks (e.g., Lins et al., 1993; Joyce et al., 2002; Toivanen et al., 2015; Mueller et al., 2016; Jia & Tyler, 2019).

Here, we used an Fp1 channel (referenced to linked mastoids) for the vertical component in all three patients and bipolar EOG2-EOG1 for patients 1 and 3 as well as F7-F8 for patient 2. Selection was based on channel availability per patient and known scalp distribution of the EOG gradient (Lins et al., 1993). To define fixation onsets, we used a probabilistic algorithm for detection of EOG events (Toivanen et al., 2015) which has been shown to achieve very high sensitivity and precision for detecting saccades and fixations - over 96% for saccade larger than 4.3 degrees of visual angle and over 90% for saccades larger or than 2.9 degrees of the visual angle; see Toivanen et al., 2015). We used an initial 20 % of data (i.e., 12 minutes) for an unsupervised training period which is required by the algorithm. The eye events were then detected in the remaining 80% of data (48 minutes) which entered further analyses. We detected 1959, 1868 and 1635 number of saccades for patients 1-3, respectively.

To sort blinks from saccades, we used a probabilistic algorithm for detecting blinks, saccades, and fixations (Toivanen et al., 2015). Briefly, there are two features of the EOG signal which are used by the classifier developed by Toivanen et al., 2015 to tell apart these three eye events. First, for fixations, the derivative of the filtered horizontal and vertical EOG is close to zero as a steady signal produces zero derivative. In turn, during saccades and eye blinks the value of the derivative is high. Hence, this feature is useful in separating fixations from blinks and saccades. To further separate fixations from eye blinks the algorithm considers a second feature which is based only on the derivative of the vertical EOG signal. The feature is defined as the difference between successive maxima and minima of the derivative, subtracted with the absolute value of the sum of these: Dv = max – min - |max+min|. The reason for this feature is that a blink produces a distinctively symmetrical pattern in the derivative of the vertical signal and should thus have a higher value for Dv than a saccade (for more details of the approach see Toivanen et al., 2015).

Broadband high frequency activity (BHA) was calculated for frequencies 80-180 Hz in 5 Hz steps using sliding Hanning tapered window (150 ms) with 6 Hz spectral smoothing. The complex-valued signal was rectified and squared to extract power and averaged across frequencies to create a single vector of BHA fluctuations. Note that we prefer using “Broadband High-frequency Activity” (BHA) rather than the common term “high-gamma”. The term “BHA” emphasizes that this signal is broadband and not necessarily an oscillation. It has distinct spectro-temporal and functional profiles as well as different generating mechanisms compared to gamma oscillation.

Continuous BHA, complex-valued time-frequency series and raw field potentials were segmented into 400 ms long non-overlapping windows relative to each fixation with 200 ms before and 200 ms after each event. For further analyses we only considered fixations that were not preceded or followed by another fixation within the +/−200ms window of interest. Based on previous studies we further constrained fixation durations to last below 800 ms (e.g., Hayhoe et al., 2003). We reasoned that longer fixations were likely spurious due to intervening small saccades that were undetected with the current setup. This resulted in 1019, 820, 637 epochs per patient 1-3, respectively. We defined epochs containing artifacts as those with gradient of field potential exceeded 5 standard deviation of the trial mean. On average (across patients) this criterion marked about 3% of epochs as containing artifacts. These segments were removed from further analyses. Phase locking value was used to quantify phase coherence in each time-frequency point. To this end, we used segmented complex-valued signal and previously described formula (Lachaux et al., 1999) with the following Matlab implementation:

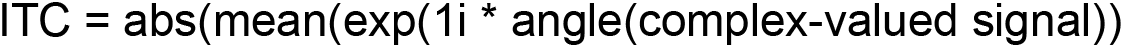

We tested whether fixation-locked field potentials, BHA, ITC or power in frequencies 1-30 Hz differed from what would be expected by chance. To test this, we created surrogate distributions (N = 1000 permutations) for each of the signals of interest (i.e., field potentials, BHA. ITC, power). The surrogate distributions for field potentials, BHA, ITC and power were created by locking intact segments of ANT signals to pseudo-events (i.e., random time points uniformly distributed across the entire recording session). For each permutation we randomly selected “pseudo-events” - these were random time points from our 1 hour long recording session which matched the number of eye events. Next, we created surrogate data segments by extracting 400 ms time epochs around each “pseudo-event”. We used the same procedure of artifact rejection to remove artifacts from our surrogate distribution as we did from our empirical data. Next, each empirical signal (fixation-onset locked field potential, PLV, BHA, power) was tested against its surrogate distribution with significance threshold set to 5^th^ and 95^th^ percentile (corresponding to p < 0.05). We used False Discovery Rate (FDR) correction for multiple comparisons across time (field potentials, BHA) and time-frequency (ITC, power; Benjamini and Hochberg, 1995).

## Acknowledgements

ML and CES supported by a Silvio O. Conte Center Grant P50 MH109429, and by R01 MH111429. TS research is funded by the European Research Council (https://erc.europa.eu/, Starting Grant 802681).

## Author Contributions

ML and CES designed the study. ML and LC collected the data. ML performed analyses. ML, CES and TS wrote the manuscript. All authors contributed to the discussion and interpretation of findings, and edited the manuscript.

